# Do stomatal movements have a limited dynamic range?

**DOI:** 10.64898/2025.12.22.695892

**Authors:** Florence Muraya, Joao Antonio Siqueira, Anne-Aliénor Véry, M. Rob G. Roelfsema

## Abstract

Stomatal movements are driven through the uptake and release of potassium (K^+^) salts by guard cells, which surround a central pore. The extrusion of K^+^ from guard cells occurs via GORK K^+^-efflux channels, but potentially the *AtKUP2, 6* and *8* encoded K^+^- transporters also play role. To test the roles of *At*KUP and GORK proteins, gas-exchange experiments were conducted with mature *Arabidopsis* leaves. These experiments revealed that loss of KUP2, 6 and 8 lowered the stomatal conductance, while it increases in the GORK loss-of function mutant. Despite the difference in stomatal conductance, the changes in transpiration induced by light and ABA had the same amplitude in wild type and mutant lines. Our data suggest that stomata have a limited dynamic range that is not affected by mutations in KUP2, 6 and 8, or GORK. We propose that these rapid stomatal movements depend on uptake and release of K^+^, inorganic and small organic anions. Consequently, changes in stomatal opening beyond the dynamic range will depend on osmolytes that cannot be rapidly released, such as larger organic anions and amino acids.

## Introduction

Transport of potassium (K^+^) plays a key role in stomatal movements, since stomatal opening depends on the accumulation of K^+^ in guard cells, while its release provokes stomatal closure (Raschke *et al*., 1988; Roelfsema & Hedrich, 2005; Kollist *et al*., 2014). Guard cells can take up K^+^ from the cell wall in a so-called “hyperpolarized state”. In this state, the plasma membrane potential is negative of -100 mV and drives K^+^ accumulation via K^+^-uptake channels. The reverse process occurs in the “depolarized state”, at membrane potentials of approximately -50 mV and allows K^+^ extrusion through K^+^ efflux channels (Thiel *et al*., 1992; Roelfsema *et al*., 2001; Hmidi *et al*., 2025).

Although the uptake, as well as the release of K^+^ is facilitated by K^+^ channels in guard cells, specific types of K^+^ channels are available for the transport in both directions. *Arabidopsis* guard cells possess K^+^ uptake channels that are encoded by several genes, all belonging to the family of cyclic nucleotide binding domain (cNBD) K^+^ channels (Jegla *et al*., 2018), which have previously been regarded as “Shaker-like channels”. The gene-family members *KAT*1, *KAT*2, *AKT*1 and *AKT*2/3 are expressed in guard cells (Szyroki *et al*., 2001), which encode K^+^ channels that activate at membrane potentials that are negative of -100 mV and thus serve as K^+^ uptake channels under normal physiological conditions. In line with multiple genes encoding these channels, stomatal opening was only inhibited if the activity of all these K^+^ uptake channels was suppressed by nonfunctional KAT2 protein, in a loss-of-function KAT2 mutant (Lebaudy *et al*., 2008).

Whereas K^+^ uptake channels are thus encoded by several genes, voltage-dependent K^+^ channels that enable K^+^ efflux are only encoded by *GORK* in guard cells (Ache *et al*., 2000; Hosy *et al*., 2003). Even though loss of GORK function causes a full loss of K^+^ efflux channels, this only results in a reduction in the speed of darkness-induced stomatal closure by a factor of 0.4. The ability of stomata of *gork* mutants to close, albeit at a reduced speed compared to wild type, suggests that guard cells possess alternative transporters that enable K^+^ efflux, but these K^+^ transport proteins have not yet been identified.

A study of Osakabe et al. (2013) suggested that a clade of KUP K^+^ transporters, comprising *At*KUP2, 6 and 8 (Very *et al*., 2014), could have a function in stomatal closure. Loss of all three genes resulted in plants with more open stomata and reduced the degree by which the drought hormone ABA can close stomata (Osakabe *et al*., 2013). Because of the potential function of the AtKUP2, 6 and 8 transporters in guard cells, we compared the stomatal responses of mature plants of triple loss-of-function *kup* mutants with wild type and *gork* mutants. These experiments revealed that loss of KUP activity reduced the stomatal conductance, while it was enhanced in the *gork* mutant. However, the magnitude of stomatal movement remained unchanged, which suggests that stomata have a fixed dynamic range for rapid stomatal movement.

## Results

### Loss of AtKUP2, 6 and 8, or GORK reduces leaf growth of mature plants

*AtKUP*2 and 6 genes are highly expressed in root tips and veins of expanding leaves (Elumalai *et al*., 2002; Osakabe *et al*., 2013), which suggests that the encoded proteins are important for the uptake and distribution of K^+^. In addition, KUP2 seems to be involved in seedling growth, since the short hypocotyl phenotype of the *shy3*- mutant is due to a point mutation in KUP2 (Elumalai *et al*., 2002).

We wondered if the three selected KUP transporters affect growth of mature plants and therefore monitored the growth of rosettes at regular intervals during a period of 6 weeks. The loss of KUP2, 6 and 8 did not cause an obvious difference in the morphology of leaves in respect to wild type, but after 31 days it limited expansion of the leaf area (Fig. 1A and B). Loss of GORK also restricted leaf growth, albeit at an earlier time point (Fig. 1A and C), and a similar reduction in growth rate was observed for the *kup6/8/gork-2* mutant (Fig. 1A and D).

**Figure 1.**
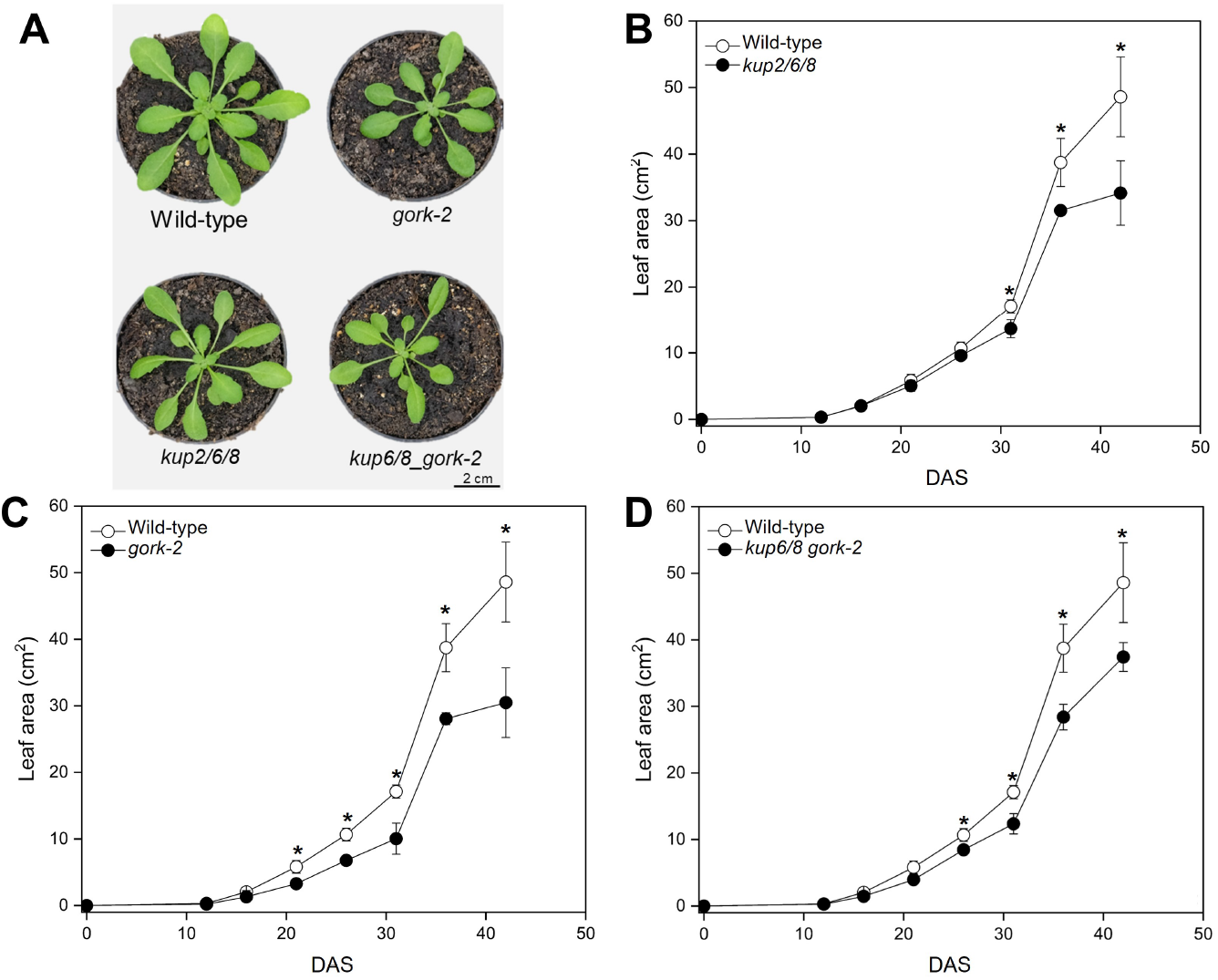
Loss of GORK and KUP2, 6 and 8 reduces leaf growth. **A**, representative images of five-week-old *Arabidopsis* plants, the genotype of plants is given below each image. Scale bar represents 2 cm. **B-D**, leaf area plotted against the growth period of 12, 16, 21, 26, 31, 36, and 42 days after sowing (DAS). Data represent mean values of 4 individuals ± standard-error (SE), open symbols show data of wild type (Col-0), while closed symbols either indicate *kup2/6/8* (B), *gork-2* (C), or *kup6/8/gork-2* (D). Asterisks indicate values that are significantly different based on 2-tailed Student’s t-test (P < 0.05).

### KUP transporters enhance the stomatal conductance but not the speed of stomatal closure

Just as *KUP6*, the *GORK* gene is highly expressed in vascular tissues and guard cells, and the expression of both genes is enhanced by drought stress (Becker *et al*., 2003; Osakabe *et al*., 2013). We therefore tested if the 2,6,8 clade of KUP transporters and GORK have a similar impact on stomatal movement. Mature leaves of *Arabidopsis* were transferred to gas-exchange cuvettes, through which air was guided at a flow speed of 1 l min^-1^.

In darkness the transpiration of *kup2/6/8* was lower than that of wild type, while that of the *gork-2* mutant was higher (Fig. 2A). The *kup6/8/gork-2* mutant showed an intermediate phenotype, as we found no significant difference in transpiration compared to wild type (Fig. 2A and B). In contrast to the results of Osakabe et al. (2013) with 3-week-old plants, the KUP2,6 and 8 transporters and GORK thus appear to have opposite impacts on the transpiration of mature plants. Application of light at a photon flux density of 125 μmol m^−2^ s^−1^ evoked stomatal opening in all lines and we found neither a difference in the magnitude, nor in the velocity of stomatal opening (Fig. 2C and D).

**Figure 2.**
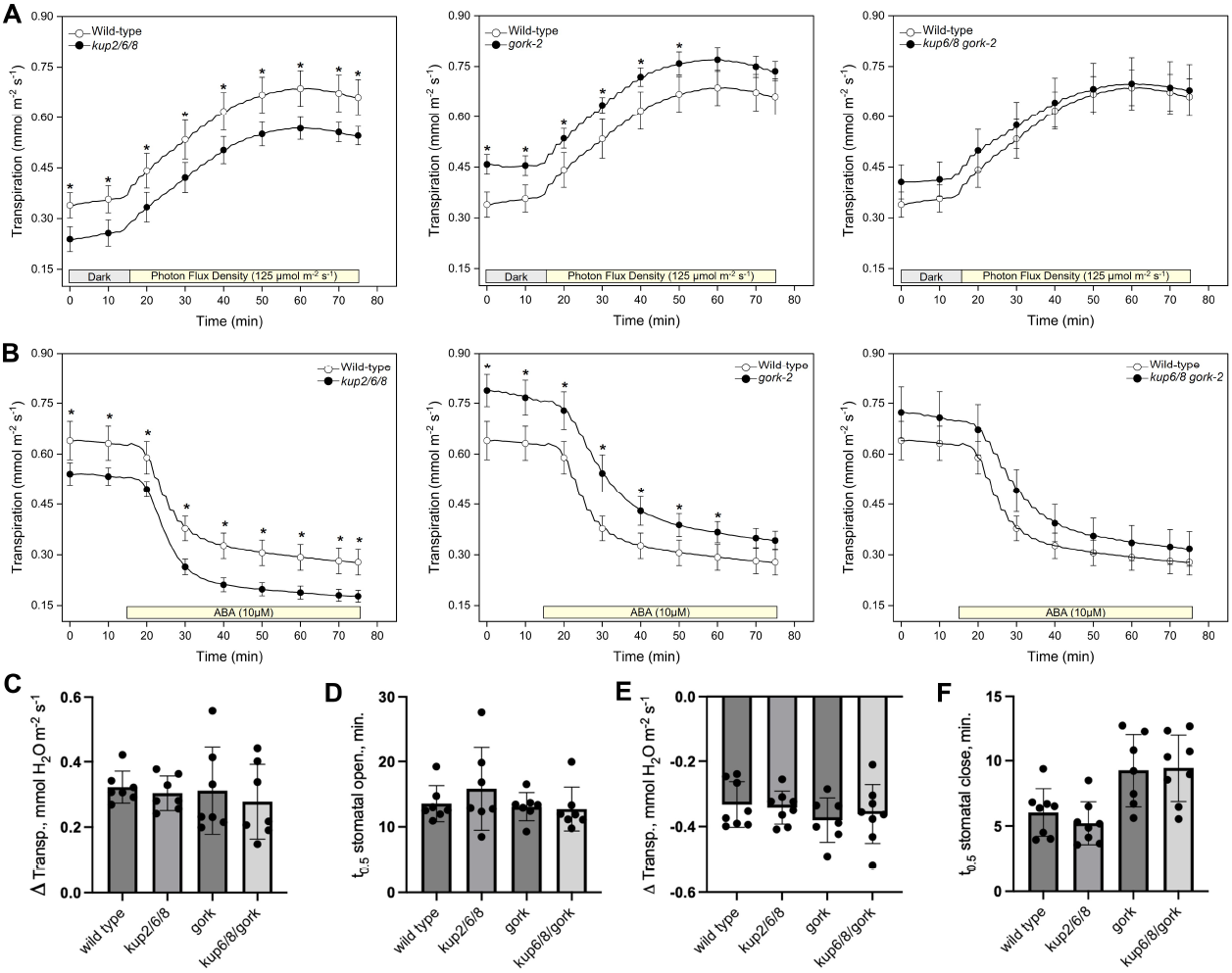
Impact of K^+^ transport proteins on leaf transpiration. **A**, Light-induced increase in leaf transpiration, measured with 3 excised leaves. The transpiration is plotted against time and light at a photon flux density of 125 μmol m^−2^ s^−1^ was applied as indicated by the bar below the traces. **B**, ABA-induced stomatal closure, measured with 3 mature leaves. The transpiration is plotted against time and ABA was supplied at a concentration of 50 µM to the petioles as indicated by the bar below the traces. **A-B** The response of wild type is shown with that of *kup2/6/8* (left panel), *gork-2* (middle panel) and *kup6/8/gork-2* (right panel). Data represent mean values of 7 experiments (except for ABA-induced closure of wild type, n=6) ± standard-error (SE). Asterisk indicates values that are different based on a 2-tailed Student’s t-test (P < 0.05). **C-F** The magnitude of change in transpiration (**C and E**), or speed of stomatal movement (**D and F**) determined from panels A and B. The speed of opening (**D**) or closure (**F**) is indicated as the half time (t_0.5_) of the maximum response, determined by non-linear regression with a single exponential function.

In addition to stomatal opening in light, we also compared the ABA-induced stomatal closure responses of wild type and the mutant lines. At the start of these experiments (photon flux density of 125 μmol m^−2^ s^−1^), the transpiration of *kup2/6/8* leaves was lower, while it was enhanced in *gork-2*, and at the same level in the *kup6/8/gork-2* line, compared to wild type (Fig. 2B). Stimulation with 50 µM ABA, provoked stomatal closure to the same extent in all lines (Fig. 2E). However, the speed of stomatal closure was approximately 2 times slower in *gork-2* and the kup6/8/gork-2 lines, in comparison to wild type (Fig. 2F).

Apparently, loss of GORK reduces the speed of stomatal closure provoked by ABA in intact leaves, just as previously shown for stomatal closure in response to darkness (Hosy et al., 2003). However, neither the loss of GORK, nor that of KUP2, 6 and 8 affects the magnitude of changes in stomatal conductance, caused by light, or ABA (Fig. 2C and E).

## Discussion

### Role of KUP transporters and GORK in long distance K^+^ transport

It is likely that the role of KUP2, 6 and 8 in plant growth is strongly influenced by conditions under which the plants are cultivated. Whereas Osakabe et al. (2013) found that loss of KUP2, 6 and 8 enhanced the leaf growth within two weeks after germination, no difference in leaf area was found in the present study. Instead, we observed similar growth rates in all lines during the first three weeks of development, and only a limitation in leaf expansion after 31 days for *kup2/6/8* and after 21 days for *gork-2*. Such a phenotype that occurs late in development would be in line with a role of the KUP and GORK proteins in long-distance K^+^ transport.

The expression of *KUP2, 6*, and *GORK* is high in the vasculature of growing leaves (Elumalai *et al*., 2002; Becker *et al*., 2003; Osakabe *et al*., 2013), which suggests that the encoded transporters contribute to K^+^ transport via the phloem, xylem, or both transport systems. Possibly GORK is required for the loading of xylem vessels in the shoot, since a homologous channel of poplar (PTORK), was found to be located at the border of vascular cells and xylem vessels (Arend *et al*., 2005). Whereas GORK channels thus may help to transport K^+^ from xylem parenchyma cells into the xylem, KUP transporters could facilitate K^+^ uptake into these cells that load the xylem (Hmidi *et al*., 2025). The collaborative role of both transporters in the vasculature would explain why their loss causes a similar reduction in growth, which occurs in a later phase of plant development.

### Opposite impact of GORK and KUP proteins on stomatal conductance

Loss of GORK appears to have a dual impact on transpiration; it reduces the speed of stomatal closure, in response to ABA and darkness, and enhances the basic stomatal conductance level (Fig. 2B and F, Fig. 3A and Hosy et al., 2003). It is likely that these features are linked to changes in the membrane potential of guard cells in the depolarized state. In this state, inhibition of K^+^ efflux channel activity shifts the guard cell membrane potentials to more depolarized values (Roelfsema & Prins, 1997; Roelfsema & Prins, 1998). As a result, loss of GORK will further depolarize guard cells and reduce the driving force for anion efflux, leading to slower speed of stomatal closure and a higher basic stomatal aperture.

**Figure 3.**
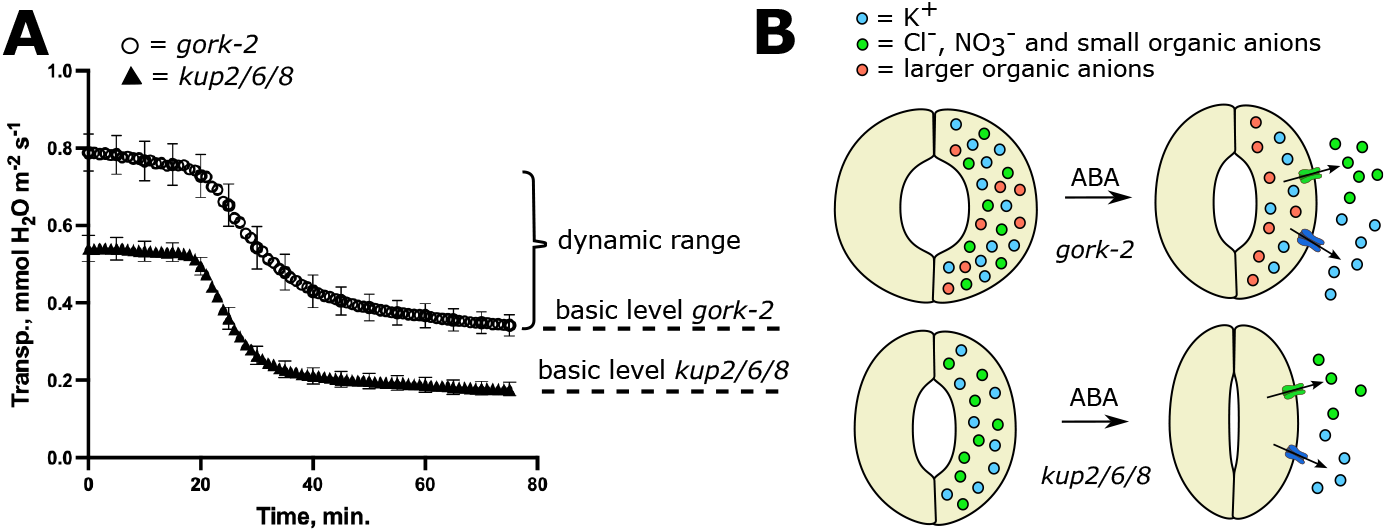
Model of basic level and dynamic range of stomatal movements. **A**, The mutations in *gork-2* and *kup2/6/8* alter the basic level of transpiration, but not the dynamic range by which stomata open and close in approximately 1 h. The average changes in transpiration, evoked by ABA, are shown for both lines (data from Fig. 2B), and the basic level and dynamic range are indicated of the right side of the traces. **B**, In *gork-2*, the stomata are further open, which is presumably due a fraction of K^+^ that is electro-neutralized by larger organic acids. These organic acids that cannot be rapidly released via S-type anion channels and ALMT12, since these channels are only permeable for inorganic anions like Cl^-^ and NO_3_^-^ and small organic acids. As a result, stomata of *gork-2* remain further open after stimulation with ABA, as their counterparts of *kup2/6/8*.

Loss of KUP2, 6 and 8 does not affect the speed of stomatal opening or closure (Fig. 2), which suggests that these transporters do not contribute to K^+^ efflux. Moreover, the mutations in these *KUP* genes lower the overall stomatal conductance (Fig. 2A and B). Apparently, GORK and KUP proteins have an opposite impact on the basic conductance of mature leaves (Fig. 3A), while the conductance is enhanced in gork-2, it is lowered in *kup2/6/8*. This suggests that KUP2, 6 and 8 increase the K^+^ levels in guard cells, probably through co-transport of K^+^ and H^+^, as was shown for the homologous transporter HAK5 (Ragel *et al*., 2015; Maierhofer *et al*., 2024).

### A limited dynamic range of stomatal movements

Despite the lower basic level in stomatal conductance of *kup2/6/8* plants than wild type, the magnitude of light- and ABA-induced stomatal movements was not altered. In the opposite way, loss of GORK enhances the overall stomatal conductance, but again, this does not affect the extend by which light and ABA provoke rapid changes in transpiration. These results strongly suggest that stomata have a fixed dynamic range, by which they can rapidly open and close (Fig. 3A). Beyond this dynamic range stomatal movements probably occur at much slower rates.

What could determine the fixed dynamic range of stomatal movements? Based on current literature, it is likely that SLAC-like (S-type) and ALMT12 channels play a major role in rapid light and ABA-induced stomatal movements (Negi *et al*., 2008; Vahisalu *et al*., 2008; Jalakas *et al*., 2021). ALMT12 is permeable for Cl^-^, and small organic acids such as malate (Meyer *et al*., 2010; Imes *et al*., 2013), while SLAC1 and SLAH3 can transport Cl^-^ and NO_3_^-^ (Geiger *et al*., 2009; Geiger *et al*., 2011). During stomatal closure, inorganic, and small organic anions, released by S-type and ALMT12 channels, thus are essential to lower the osmotic content of guard cells.

Guard cells can only rapidly release as much K^+^, as is electrically neutralized by Cl^-^, NO_3_^-^ and small organic anions, that are extruded via S-type and ALMT12 channels. As a result, this fraction of K^+^ and anions represents the mobile pool of osmolytes (Fig. 3). This major role of anion channels is supported by results with *slac1 and amlt12* mutants (Merilo *et al*., 2013; Jalakas *et al*., 2021). Gas exchange experiments with these mutants showed that loss of SLAC1 and ALMT12 not only enhances the basic level of stomatal conductance but also alters the dynamic range. In contrast, the KUP and GORK K^+^ transporters affect the basic conductance, but not the dynamic range of stomatal movement.

The fraction of K^+^ that is neutralized by larger organic anions, will have a low mobility and therefore does not contribute to rapid stomatal movements (Fig. 3B). Instead, these larger organic anions may be slowly metabolized or released from guard cells at low flux rates. Consequently, the accumulation of K^+^ and larger organic anions by guard cells will increase the basic level in stomatal conductance but not affect the dynamic changes (Fig. 3).

### Regulation of KUPs by ABA

The enhanced level of stomatal conductance in wild type, relative to *kup2/6/8*, suggests that KUP transporters enhance K^+^ uptake, which is electrically neutralized by organic anions with low mobility. As a result, wild type plants transpire at a higher rate than *kup2/6/8*, which cannot be reduced by activation of S-type and ALMT12 anion channels. The incomplete closure of stomata is likely to be of disadvantage during drought, as plants will keep losing water via these pores. During dry periods, ABA may therefore inhibit KUP transporters, which would slowly reduce the stomatal aperture. This response may be involve OST1 (also known as SnRKE, or SnRK2.6), which was shown to phosphorylate the C-terminal of KUP6 and 8 (Osakabe *et al*., 2013). ABA thus may provoke an OST1-dependent slow reduction of the basic stomatal conductance level, in addition to rapid stomata closure by activation of S-type and ALMT12 anion channels (Roelfsema *et al*., 2004; Geiger *et al*., 2009; Imes *et al*., 2013).

## Conclusion

Our study indicates that stomata have a limited dynamic range for fast movements, which occur in approximately an hour after stimulation with light, or ABA. Loss-of function mutations in GORK and KUP proteins do not affect the dynamic range of stomatal movements but only modify the basic level of stomatal conductance. It is likely that the dynamic stomatal movements are mainly due to guard cell uptake of Cl^-^ and NO_3_^-^, and the release of these inorganic anions via S-type anion channels. In addition to these fast stomatal movements, light, CO_2_ and ABA probably also cause slow changes in stomatal movement, due to modulation of the basic level of stomatal conductance opening, which is linked to accumulation of low-mobile organic anions. Future studies should address the importance of these slow stomatal movements for the adaption of plants to drought and other stress conditions.

## Material and methods

### Plant material, growth conditions and analysis

All lines of *Arabidopsis thaliana* used in this study were in the Col-0 background and obtained from the RIKEN BioResource Research Center, Tsukuba, Ibaraki, Japan. The following mutant lines were used; psi00288 (*kup2/6/8*), psi00289 (*kup6/8/gork-2*) and psi00290 (*gork-2*). The plants were cultivated in a growth cabinet with 60% relative humidity, a cycle of 12 h light: 12 h dark, temperatures of 22°C (light) and 18°C (dark), and a photon flux density of 100 μmol m^−2^ s^−1^. After 12 days, the seedlings were transferred to pots (diameter 6 cm) and grown up to 6 weeks at the same conditions as described above.

### Gas exchange measurements and analysis

Gas exchange measurements were conducted with three excised mature leaves, which were placed with their petiole through a hole (diameter 5 mm) in a vial (volume 10 ml). The transpiration was monitored in a custom-made gas exchange setup (Muller *et al*., 2017), equipped with two infrared gas analyzers (IRGA; LI 7000; Li-Cor, Lincoln, NE, USA). The flux of air inside the cuvettes was adjusted by mass flow meters (red-y; Vögtlin, Muttenz, Switzerland), to a flow of 1 l min^−1^, 50% relative humidity and 400 ppm CO_2_. White light was provided by LEDs (CXA2520-0000-000N0YK427F; Cree, Durham, NC, USA), at a photon flux density of 125 μmol m^−2^ s^−1^. Experiments were conducted with three mature leaves, During the measurement, ABA was applied to the vial at a final concentration of 50 μM.

The rate of stomatal opening, or closure, was determined by non-linear regression of transpiration data from the time point of light, or ABA application, for opening and closure respectively. Single experiments were analysed using a single exponential function in Graphpad Prism (Graphpad, Boston, MA, USA). The magnitude and speed of stomatal movement are presented as single data points for each measurement, the average values and standard deviation.

## Acknowledgements

We thank Peter Ache, Anne Baerwolff and Heike Müller (University of Würzburg, Germany) for assistance with the gas-exchange experiments.

